# Semantic composition in experimental and naturalistic paradigms

**DOI:** 10.1101/2023.10.31.564951

**Authors:** Jixing Li, Marco Lai, Liina Pylkkänen

**Affiliations:** City University of Hong Kong, 83 Tat Chee Ave, Kowloon Tong, Hong Kong; New York University

## Abstract

Naturalistic paradigms using movies or audiobooks have become increasingly popular in cognitive neuroscience, but connecting them to findings from controlled experiments remains rare. Here, we aim to bridge this gap in the context of semantic composition in language processing, which is typically examined using a “minimal” two-word paradigm. Using magnetoencephalography (MEG), we investigated whether the neural signatures of semantic composition observed in an auditory two-word paradigm can extend to naturalistic story listening, and vice versa. Our results demonstrate consistent differentiation between phrases and single nouns in the left anterior and middle temporal lobe, regardless of the context. Notably, this distinction emerged later during naturalistic listening. Yet this latency difference disappeared when accounting for various factors in the naturalistic data, such as prosody, word rate, word frequency, surprisal, and emotional content. These findings suggest the presence of a unified compositional process underlying both isolated and connected speech comprehension.

## Introduction

Naturalistic paradigms utilizing movies or audiobooks have gained considerable popularity in the field of cognitive neuroscience. Within the realm of language studies, these approaches provide valuable insights into language processing in real-world contexts, allowing for the examination of a broader range of linguistic phenomena (Alday, 2019; Brennan, 2016; Kandylaki & Bornkessel-Schlesewsky, 2019). During the comprehension of narratives, linguistic processes unfold naturally across multiple levels, including words, phrases, sentences, and discourse, each operating on distinct timescales. Computational models have often been employed to isolate these sub-processes and target specific linguistic levels (Brennan et al., 2016; Brennan & Pylkkanen, 2012; Goldstein et al., 2022; Huth et al., 2016; Schrimpf et al., 2021; Wehbe et al., 2021). For instance, relevant neural signals for semantic features of words were identified using word embedding models (Huth et al., 2016). Nevertheless, it is important to recognize that narrative comprehension involves a multitude of processes beyond the domain of language, including attention, emotion, social-cognitive functions, and memory encoding and retrieval (Hasson & Egidi, 2015). Consequently, it is possible to misattribute regions involved in these non-linguistic processes as core language regions.

Controlled experiments, on the other hand, are designed to isolate relevant cognitive processes by comparing conditions that differ solely in the component of interest. Early neurolinguistic experiments typically compared sentences with simple and complex syntactic structures, such as center-embedded and object-relative clauses (Stromswold et al., 1996), garden-path sentences (Bever, 1970), or implausible sentence completions (Kutas & Hillyard, 1980). This work was later complemented by research on basic meaning composition using a “minimal” two-word paradigm, where compositional phrases such as “red boat” were contrasted with single nouns such as “xkq boat” (Bemis & Pylkkanen, 2011; Bemis & Pylkkänen, 2013; reviewed in Pylkkänen, 2019). The underlying rationale behind these experimental manipulations is based on the concept of subtraction, although this approach has faced criticism as the brain is unlikely to behave like a linear system (Friston et al., 1996). Moreover, the experimental stimuli often diverge from everyday language use (Brennan, 2016). Thus, while controlled experiments have been widely embraced in neurolinguistics, their applicability to language processing in real-world contexts remains uncertain.

To compare the findings from experimental and naturalistic paradigms, we conducted a study incorporating both designs, with a specific focus on meaning composition, a fundamental function underlying human language’s expressive capacity. The left anterior temporal lobe (LATL) has been consistently implicated in the effects of semantic composition, as demonstrated in studies using a two-word design (e.g., Blanco-Elorrieta et al., 2018; Li & Pylkkänen, 2021; Westerlund & Pylkkänen, 2014, reviewed in Pylkkänen, 2019). However, the generalizability of these findings to naturalistic settings has received limited exploration. Here, we trained feed-forward neural network (FFNN) classifiers to differentiate between MEG source estimates for adjective-noun phrases and single nouns, both in the two-word (e.g., “green glass” vs. “glass”) and naturalistic settings (e.g., “…soft music…” vs. “…a bath…”). To examine the generalizability of the classifiers, we tested the classifiers trained in the experimental setting on the naturalistic data using the temporal generalization method (TGM; King & Dehaene, 2014), and vice versa. Our results revealed that the left anterior and middle temporal lobe consistently differentiated between phrases and single nouns in both the experimental and naturalistic contexts, aligning with previous findings concerning semantic composition (see Pylkkänen, 2019 for a review).

Notably, the combinatory effect occurred much later in the naturalistic setting, be attributed to additional processing demands imposed by other information present in the naturalistic data, such as prosody, word rate, word frequency, surprisal of incoming words, and emotional content. To examine this possibility, we conducted further analyses by regressing out these effects and re-evaluating the classification results. The revised analyses revealed an earlier composition effect in the naturalistic setting, closely resembling the pattern observed in the two-word setting. These findings provide compelling evidence for a unified compositional process underlying both the experimental and naturalistic contexts, once the confounding effects are accounted for.

## Materials and Methods

### Experimental design

The MEG experiment consists of a two-word session and a naturalistic listening session and was presented within a larger protocol that also included production tasks. Fitting multiple tasks into a single recording session manageable for children was a major design constraint. While most of the prior comprehension literature has used reading, the current study was auditory, as we wanted the paradigm to be suitable even for children who cannot read yet. Reading and listening were contrasted in Bemis & Pylkkänen (2013) who did observe a LATL sensitivity to a composition effect for both reading and listening.

In the two-word session, participants listened to both adjective-noun phrases (e.g., “green glass”) and single nouns that were preceded by a non-lexical “mmm” sound, chosen for naturalness in a speech context (“mmm glass”). After the auditory stimulus, participants selected a matching picture from a set of 8 pictures. This task differed from the prior minimal composition studies which have only used one matching or mismatching task picture (Bemis & Pylkkanen, 2011). The reason for our larger set of pictures was that this decreased the chance of an accurate response by chance, making the behavioral data more informative if the task were to be used in, say, a clinical setting. There were 6 unique color words (“red, pink, blue, green, black, white”) and 6 unique nouns (“glass, comb, door, sword, heart, house”), and they were randomly combined to form adjective-noun phrases. Each participant received a unique randomisation. A total of 50 phrases and 50 single nouns were presented. Some adjective-noun combinations were presented more than once, and each noun was repeated 8-9 times.

### Participants

Participants were 20 healthy adults (15 females, *M*=27.8 years, SD=13.2) and 11 school-age children (6 females, *M*=9.4 years, SD=2.3) with normal hearing and normal or corrected-to-normal vision. We included children in our sample to test whether language proficiency and the development of cognitive functions such as social and emotional functions may affect language processing in natural and unnatural contexts. The sample size of children is relatively small due to the pandemic. All of the participants were strictly qualified as right-handed on the Edinburgh handedness inventory (Oldfield, 1971). They self-identified as native English speakers and gave their written informed consent prior to participation, in accordance with New York University.

### Experiment procedures

Before recording, each subject’s head shape was digitized using a Polhemus dual source handheld FastSCAN laser scanner. Participants then completed the experiment while lying supine in a dimly lit, magnetically shielded room (MSR). MEG data were collected using a whole-head 156-channel axial gradiometer system (Kanazawa Institute of Technology, Kanazawa, Japan). The two words were presented for 875 ms each, and an image with 8 objects appeared on screen 600 ms after the second word. Subjects then selected the correct object that matched the auditory stimuli. No feedback was provided. The inter-stimulus interval was normally distributed with a mean of 300 ms (SD=100 ms). Order of stimulus presentation was randomized and each participant received a unique randomisation. After the two-word session, participants completed a naturalistic listening session where they passively listened to an audio excerpt consisted of 4 stories from the YouTube channel “SciShow Kids”. The two-word session lasted around 20 minutes and the naturalistic listening session lasted about 12 minutes. After the MEG recording, participants completed 4 picture-matching questions on the contents of the stories (See Figure 1A for the experiment procedure).

**Figure 1.**
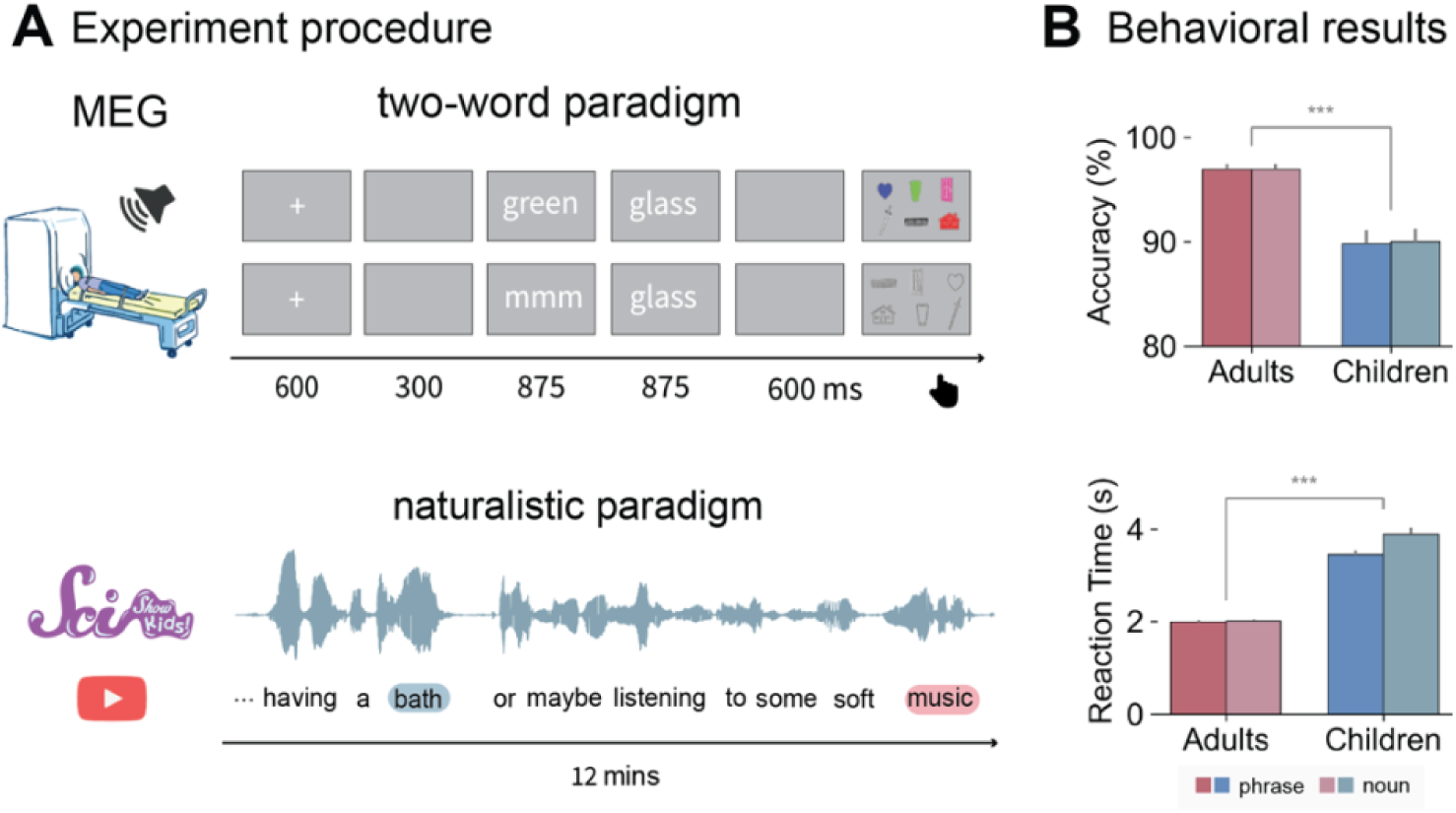
Experimental design and behavioral results. **A.** Experimental design and trial structure. In the two-word session, participants selected a picture from 8 pictures that matched the preceding words in each trial. Half of the target pictures matched and half did not. Activities recorded from the onset of the second word to 875 ms after the second word were analyzed. In the naturalistic listening session, participants passively listened to a 12-min audio excerpt from the YouTube channel “SciShow Kids”. Participants completed a picture-matching task after the listening session to ensure comprehension. **B.** Behavioral results on the two-word task. Mean predicted accuracy and reaction time for the phrase and noun conditions across the adults and children group. A two-way mixed ANOVA revealed significant differences between the groups in both accuracy (p=.002) and reaction time (p<.001). Composition was not significant for either accuracy or reaction time for either group. Error bars indicate 1 standard error.

### MEG data acquisition and pre-processing

MEG data were recorded continuously at a sampling rate of 1000 Hz with an online 0.1 to 200 Hz band-pass filter. The raw data were first noise reduced via the Continuously Adjusted Least-Squares Method (Adachi et al., 2001) and low-pass filtered at 40 Hz. Independent component analysis (ICA) was then applied to remove artifacts such as eye blinks, heartbeats, movements, and well-characterized external noise sources. MEG data from the two-word task were segmented into epochs spanning 100 ms pre-stimulus onset to 1750 ms post-stimulus onset. MEG data from the naturalistic task were segmented into epochs from the onset to 875 ms after the target word. The target words include words at the boundary of single nouns and adjective-noun phrases in the naturalistic stimuli. Single nouns and adjective-noun phrases were annotated based on the Stanford part-of-speech tagger (Toutanova et al., 2003).

Epochs containing amplitudes greater than an absolute threshold of 2000 fT were automatically removed. Additional artifact rejection was performed through manual inspection of the data, removing trials that were contaminated with movement artifacts or extraneous noise. The whole epoch rejection procedure results in an average rejection rate of 7.6% (SD=5.1%) for the adult participants and an average rejection rate of 11.1% (SD=5%) for the child participants.

We then computed the cortically constrained minimum-norm estimates (Hämäläinen & Ilmoniemi, 1994) for each epoch for each participant. To perform source localization, the location of the participant’s head was first coregistered with respect to the sensor array in the MEG helmet. We used the FreeSurfer (http://surfer.nmr.mgh.harvard.edu/) “fsaverage” brain with rotation and translation and then scaling the average brain to match the size of the head scan. A source space of 2562 source points per hemisphere was generated on the cortical surface for each participant. The Boundary Element Method (BEM) was employed to compute a forward solution, explaining the contribution of activity at each source to the magnetic flux at the sensors. For the two-word data, channel-noise covariance was estimated based on the 100 ms intervals prior to each artifact-free trial. The naturalistic data were baseline-corrected using the mean of the whole epoch. The inverse solution was computed from the forward solution and the grand average activity across conditions. To lift the restriction on the orientation of the dipoles, the inverse solution was computed with “free” orientation, meaning that the inverse operator places three orthogonal dipoles at each location defined by the source space. When computing the source estimate, only activity from the dipoles perpendicular to the cortex was included. For each trial within one condition, the same inverse operator for that condition was applied to yield dynamic statistical parameter maps (dSPM) units (Dale et al., 1999) using an SNR value of 3. The final source estimates were decimated by a factor of 5 to save computing time. All data preprocessing steps were performed using MNE-python (v.0.24.0; (Gramfort et al., 2014)).

### Behavioral data analyses

Accuracies were analyzed using a generalized linear mixed-effects model (GLMM) with binomial error distribution, and the log-transformed RTs were analyzed using a linear mixed-effects model. Our fixed effects include the binary variables Composition (single nouns vs. phrases) and Age (adults vs. children). Subject-level variability was included as random intercepts. The GLMM analyses were conducted via the “lme4” package (Bates et al., 2015) in R (v4.2.1) and RStudio (v022.12.0+353). The statistical significance of fixed effects was estimated using the “lmerTest” package (Kuznetsova et al., 2017), in which Satterthwaite’s approximation was applied to estimate degrees of freedom (see Figure 1B for the results).

### Phrasal and noun representations in LLMs

To gain insights into the neural representations of phrases and single nouns in the two-word and naturalistic contexts, we first examined phrasal and noun representations in isolated two words and narratives in a large language model (LLM). Recent LLMs have achieved extraordinary performance in language comprehension tasks and have been suggested to share some computational principles with the human brain (e.g., Caucheteux & King, 2022; Goldstein et al., 2022; Schrimpf et al., 2021). Here, we first extracted each layer’s embeddings from the pre-trained GPT2-large model (Radford et al., 2019) for the nouns in single nouns and adjective-noun phrases in the two-word (e.g., “green glass” vs. “glass”) and narrative contexts (e.g., “…soft music…” vs. “…a bath…”). We then applied multidimensional scaling (MDS), a dimensionality reduction technique to visualize the last layer’s embedding of each adjective-noun phrase and single noun in the two-word and naturalistic contexts to 2 dimensions (see Figure 2A). We also computed the cosine distance between each layer’s embeddings for single nouns and adjective-noun embeddings (see Figure 2B). The pretrained GPT2-large model was obtained from the transformers (v4.10.2) package in python.

**Figure 2.**
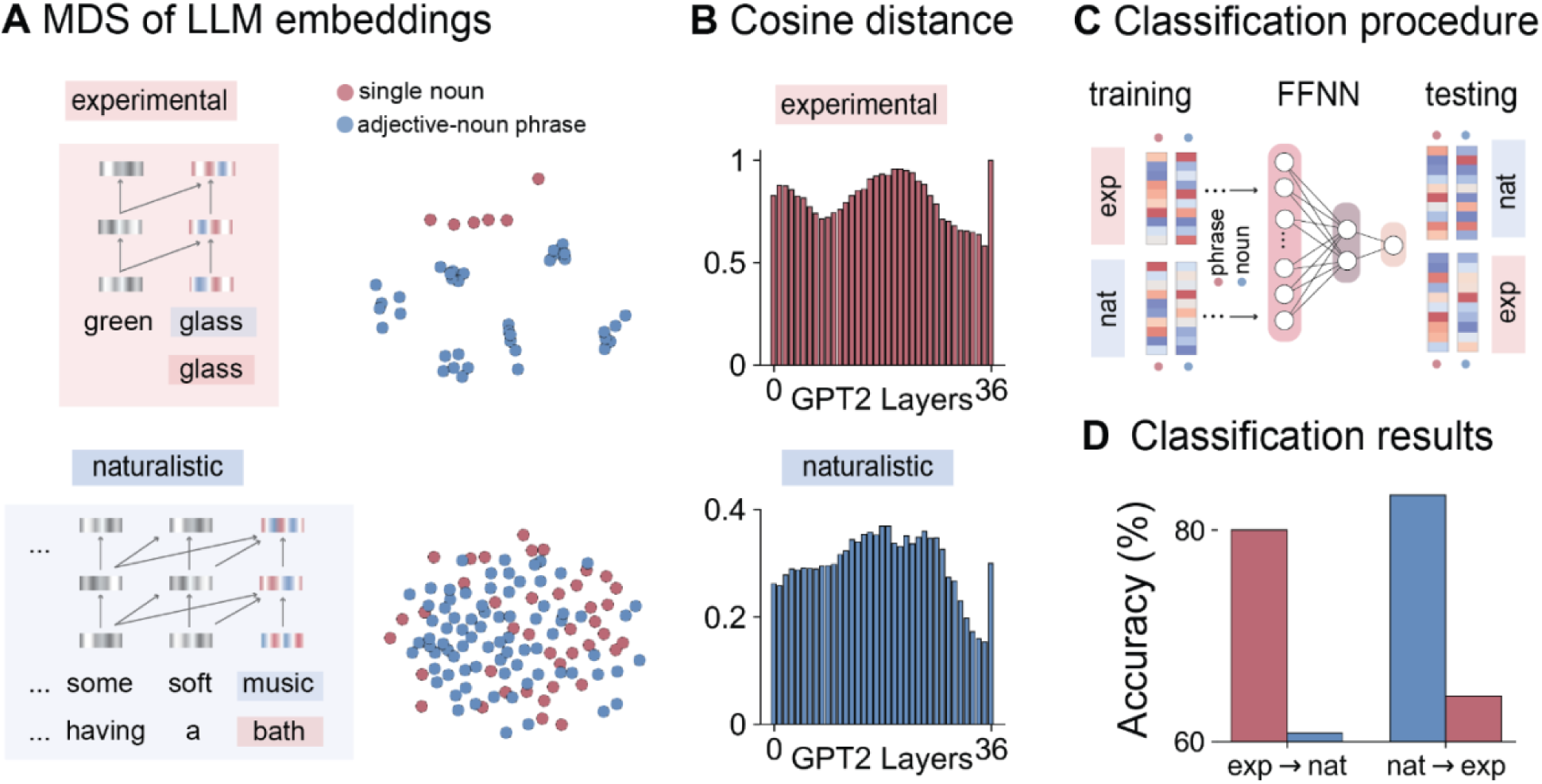
Phrasal and noun representations in two-word and naturalistic contexts in an LLM. **A.** Multidimensional scaling (MDS) of the last layer’s embeddings of adjective-noun phrases and single nouns in two-word and naturalistic contexts in GPT-2. **B.** Cosine distance of each layer’s embeddings of adjective-noun phrases and single nouns in two-word and naturalistic contexts in GPT-2. **C.** A feed-forward neural network classifier was trained to distinguish the last layer’s embeddings of nouns in single nouns and adjective-noun phrases in the two-word context, and tested on the last layers of nouns in single nouns and adjective-noun phrases in the naturalistic context. Conversely, a classifier was trained on the naturalistic context and tested on the two-word context. **D.** Classification results on the LLM’s embeddings. The classifier trained in the two-word context achieved an accuracy of 80% in distinguishing phrases from nouns and an accuracy of 60.8% when applied to the naturalistic context. The classifier trained in the naturalistic context achieved an accuracy of 83.3% and an accuracy of 64.3% when tested in the experimental context.

### Classification on LLM embeddings for phrases and nouns

We trained a feed-forward neural network (FFNN) classifier to distinguish the nouns in single nouns and adjective-noun phrases using the two-word stimuli and tested the classifier on the nouns in single nouns and adjective-noun phrases in the naturalistic text. Adjective-noun phrases were annotated using the Stanford part-of-speech tagger (Toutanova et al., 2003) and were manually checked. Conversely, we also trained an FFNN classifier on the naturalistic data and tested it on the two-word data. The FFNN contains one hidden layer with two units (see Figure 2C). To control for the confounding factor that the nouns in single nouns were the initial token whereas the nouns in adjective-noun phrases were not, we performed a linear regression model using the binary variable “word position” to predict each layer’s embeddings. We took the residuals of the model for the classification analyses. The classification analyses were performed using the python package scikit-learn (v0.22.1).

### Searchlight multivariate pattern classification on MEG data

We conducted searchlight multivariate pattern classification analyses on the source-localized MEG data within a left-lateralized language mask for each subject. The language mask (see the pink region in Figure 3A) covered regions including the whole left temporal lobe, the left inferior frontal gyrus (LIFG; defined as the combination of BAs 44 and 45), the left ventromedial prefrontal cortex (LvmPFC; defined as BA11), the left angular gyrus (LAG; defined as BA39) and the left supramarginal gyrus (LSMA; defined as BA 40). The left AG and vmPFC have also been implicated in previous literature on conceptual combination (Bemis & Pylkkanen, 2011; Price et al., 2015) and the LIFG and the LMTG have been suggested to underlie syntactic combination (Flick & Pylkkänen, 2020; Hagoort, 2005; Lyu et al., 2019; Matchin et al., 2019; Matchin & Hickok, 2020).

**Figure 3.**
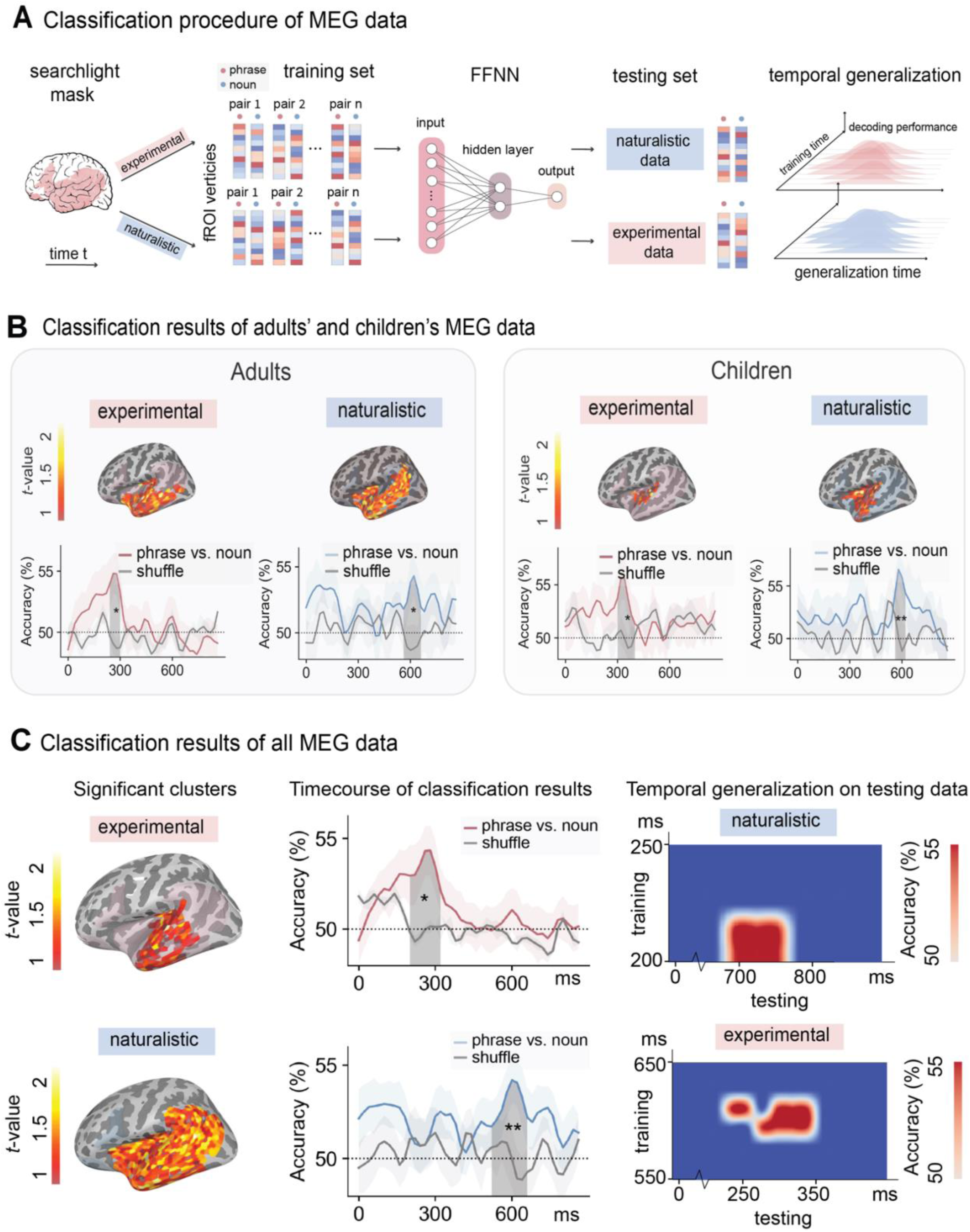
Classification analyses procedure and results on the MEG data. **A.** Following the classification analysis of phrases vs. nouns in LLMs, we trained feed-forward neural network (FFNN) classifiers to distinguish phrases from nouns in one context and tested in another context. The same classification was applied independently with a searchlight radius of 20 sources within a language mask and at every timepoint. Classification accuracies for the training data were averaged over subjects at source and time point minus the chance level of 50% was submitted to a one-sample *t*-test and the statistical significance was determined by a TFCE correction with 10,000 permutations. At the testing time, we applied the temporal generalization method and tested the classifiers’ performance at each time point on every timepoint in the testing data. **B.** Classification results of adults’ and children’s MEG data. For adults, the classifiers trained on the experimental data can distinguish phrases from single nouns in the left anterior and middle temporal lobe from 240-320 ms (*p*=0.025) after the onset of the target word. The classifiers trained on the naturalistic data can distinguish phrases from single nouns in the similar left anterior and middle temporal regions from 560-680 ms (*p*=0.03) after the onset of the target word. For children, the classifiers trained on the experimental data can distinguish phrases from single nouns in the left middle temporal lobe from 300-420 ms (*p*=0.014) after the onset of the target word. The classifiers trained on the naturalistic data can distinguish phrases from single nouns in the left anterior and middle temporal regions from 560-640 ms (*p*=0.001) after the onset of the target word. **C.** Classification results of all MEG data. When trained on experimental data, the classifiers can distinguish phrases from single nouns in the left anterior and middle temporal lobe from 200-340 ms (*p*=0.005) after the onset of the word. When tested on the naturalistic data, the TGM results suggest that the classifiers from 200-220 ms in the training data can significantly distinguish phrases from nouns from 700-760 ms in the testing data. When trained on the naturalistic data, the classifiers can distinguish phrases from single nouns in the whole left temporal lobe from 520-680 ms (*p*=0.001) after the onset of the word. When tested on the experimental data using TGM, the classifiers from 620-640 ms in the training data can significantly distinguish phrases from nouns from 220-340 ms in the testing data. The grey lines represented shuffled classification results.

We trained feedforward neural network (FFNN) classifiers to pairwise combinations of the MEG data for single nouns and phrases in the two-word and naturalistic experiments (see Figure 3A).

The FFNN contains one hidden layer with two units. The binary classifiers were separately applied to all spatiotemporal timepoints, with a radius of 20 sources. The same analysis pipeline was applied to each subject. At the group level, the classification accuracy averaged over subjects at each timepoint minus the chance level of 50% was submitted to a one-sample one-tailed t-test with threshold-free cluster enhancement (TFCE) correction (Smith & Nichols, 2009) for 10,000 permutations (see the first two columns of Figure 3B for the results). The analysis time window was between 0-875 ms after the onset of the second word.

### Testing the classifiers using the temporal generalization method (TGM)

The classifiers trained to distinguish the MEG data for single nouns and phrases in the two-word task were tested on the MEG data for single nouns and phrases in the naturalistic task using TGM. TGM allows us to probe compositional processing in the brain over time by training the classifier using data from one time period and testing the classifier on data from all time periods. This method is particularly useful for neuroimaging data with high temporal resolution (e.g., EEG, MEG), and it has been successfully applied in other domains of cognitive neuroscience such as memory (Meyers, 2018), vision (Dobs et al., 2019), audition (King et al., 2014), etc. The results of TGM is a 2D matrix, where the color at point *i, j* indicated prediction accuracy when the model is trained using data at time *i* and tested with data at time *j* (see Figure 3A for the classification procedure).

Similarly, the classifiers trained on the naturalistic data were tested on the experimental data using TGM. During testing, each classifier trained from training data at a timepoint was applied to testing data at all timepoints. This procedure led to 2 TGM matrices of classification performance, one for training on experimental data and testing on naturalistic data, and one for training on naturalistic data and testing on experimental data. Statistical significance is decided based on a cluster-based one-sample one-tailed *t*-test with 10,000 permutations (Maris & Oostenveld, 2007), comparing the 2D matrix to a chance level of 0.5 (see the last column of Figure 3B for the results). The classification analyses were performed using the python package scikit-learn (v0.22.1) and the statistical analyses were performed using the python package eelbrain (v0.38).

### MDS of MEG data of phrases and nouns

We extracted the MEG data from the significant clusters derived from the classification analyses (see the first column in Figure 3B). We then applied MDS to the MEG source estimates of each target word in the two-word and naturalistic contexts. We also plotted the temporal dynamics of the 2D representations of the single-nouns and adjective-noun phrases in the “experimental” and “naturalistic” state space (see Figure 4).

**Figure 4.**
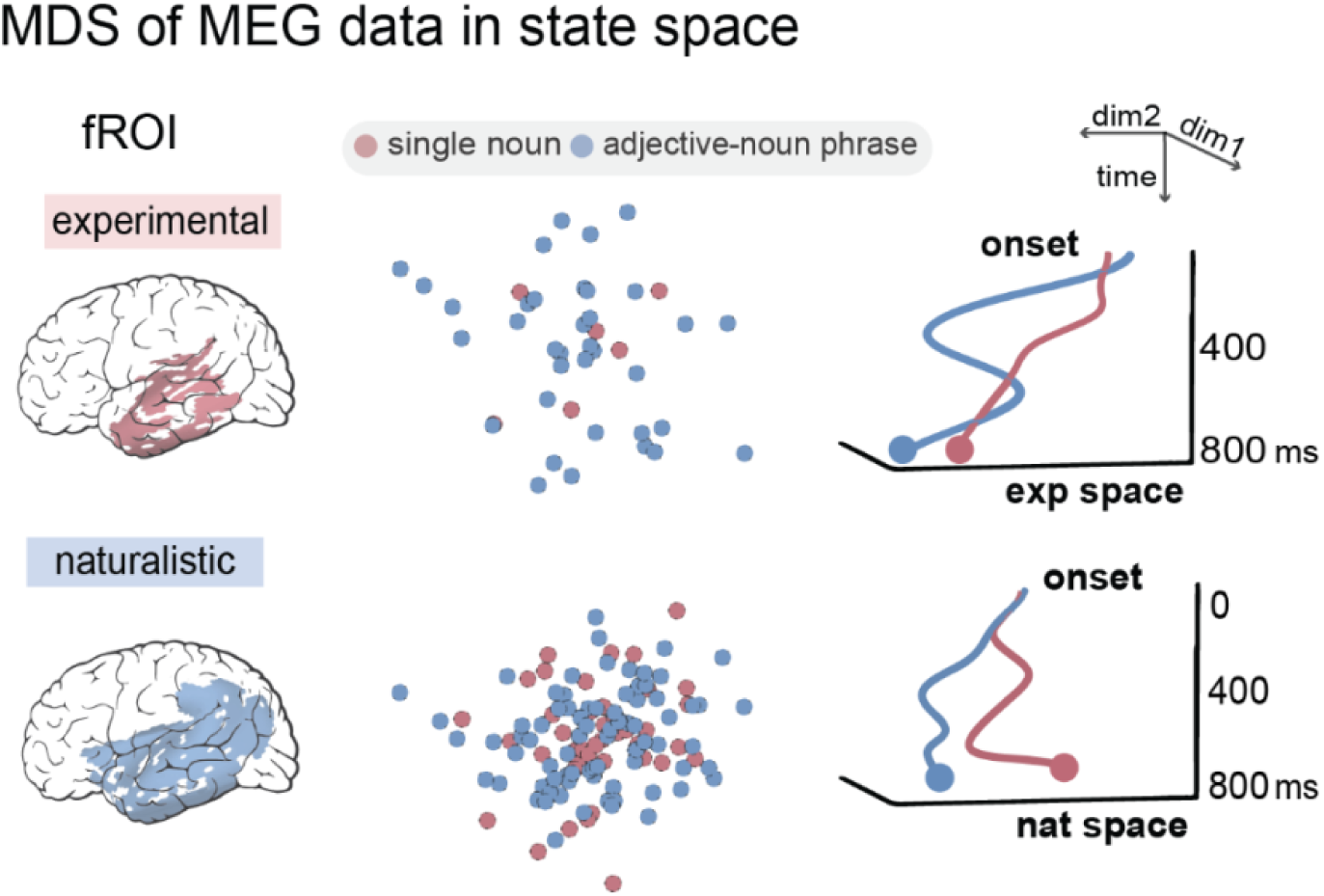
MDS of the neural representations for phrases and nouns in the experimental and naturalistic contexts. The MEG source estimates from the significant spatiotemporal clusters in the classification analyses were extracted and reduced to two dimensions using MDS. The timecourses of the MDS representations of phrases and nouns in the experimental state space suggested larger distance in an earlier time window. For the naturalistic data, the timecourses of the MDS representations diverted from the middle to the end of the whole epoch.

### Multiple regression on the naturalistic MEG data

Naturalistic stimuli differ from two-word stimuli in many dimensions. For example, the stimuli in the two-word task had lower surprisal as they were repeatedly presented during the experiment. Surprisal evoked by an incoming word indicates the amount of information that was not predictable from the context (Hale, 2001; Levy, 2008), and is calculated as the negative logarithm of the probability assigned to the actual next word. A slower presentation rate of words (875 ms) in the two-word task may also facilitate faster composition compared to words that are much faster during naturalistic speaking. Other linguistic factors such as richer prosodic information and different word frequency may also induce additional processes that delayed the composition effect. In addition, processes beyond the language domain may be involved during narrative understanding. Emotional arousal and valence, for example, have been shown to also evoke activity in the language network (Wallentin et al., 2011).

To understand whether these factors that are underlying the “naturalness” of the narrative stimuli contributed to the late composition effect, we conducted a multiple regression model to regress out these factors (see Figure 5A). Our dependent variable is the source estimates of each subject’s naturalistic data. Our regressors included the peak intensity and f0 of the target words, word rate, word frequency, word surprisal based on the GPT-2 language model (Radford et al., 2019), emotional valence and arousal indicated by human judgment on Amazon Mechanical Turk (see details of the regressors below). Both the dependent and independent variables were z-scored. Pearson’s *r* correlations among the regressors were examined to ensure no collinearity among the regressors (see Figure 5C).

**Figure 5.**
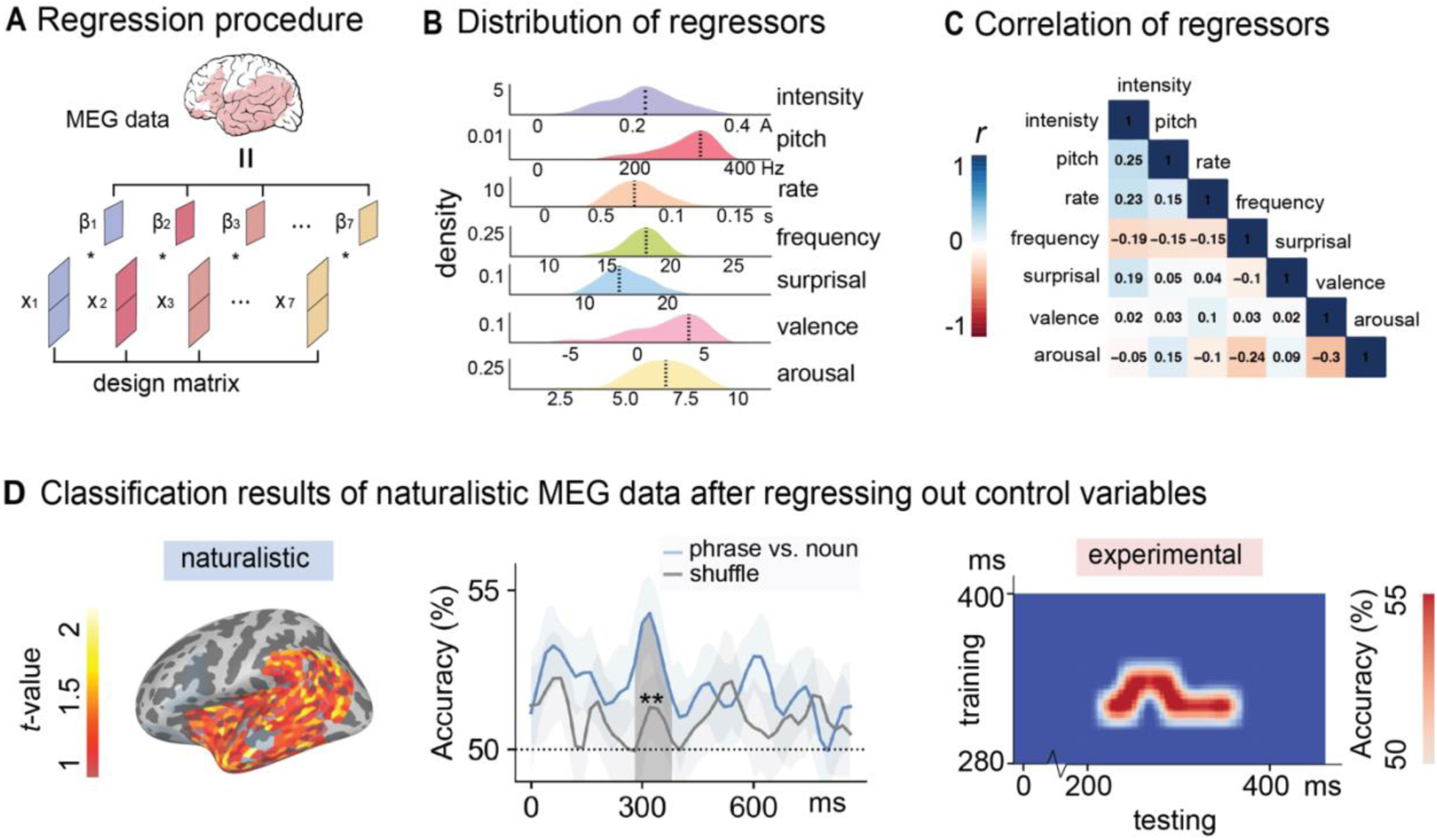
Regression analyses procedure and the classification results of the naturalistic data after regressing out control variables. **A.** We applied a linear regression model to predict the source estimates of the target words in the naturalistic data for each subject. **B.** Distribution of the regressors. Our regressors include intensity and pitch for the audio, word frequency, presentation rate, surprisal of the word given previous context, and emotional arousal and emotional valence of the text. **C.** Correlations among the regressors. The correlation matrix suggested low correlation among the predictors. **D.** Classification results after controlling for the regressors. We performed the same classification analyses on the residuals of the source estimates after regression. When trained on the naturalistic data, the classifiers can distinguish phrases from single nouns in a large cluster in the left temporal lobe from 280-400 ms (*p*=0.002) after the onset of the word. When tested on the experimental data using TGM, the classifiers from 320-340 ms in the training data can significantly distinguish phrases from nouns from 220-360 ms in the testing data. The grey lines represented shuffled classification results.

### Intensity and pitch

Root mean square (RMS) intensity and the fundamental frequency (f0) for every 10 ms of the audio were extracted using the Voicebox toolbox (http://www.ee.ic.ac.uk/hp/staff/dmb/voicebox/voicebox.html). Peak RMS intensity and peak f0 within the during of each word in the naturalistic stimuli were used to represent the intensity and pitch information for each word.

### Word rate

Since word duration is largely determined by the length of the word, we computed the presentation rate of each word as the duration of each word in milliseconds divided by the number of letters in the word. A slow presentation rate indicates words with longer duration and fewer letters, while a fast presentation rate suggests a shorter presentation of long words.

### Word frequency

Log-transformed unigram frequency of each word in the naturalistic stimuli was estimated using Google Books Ngram Viewer, Version 2012070129 (http://storage.googleapis.co m/books/ngrams/books/datasetsv2.html).

### Surprisal

The predictability of each word in the naturalistic stimuli given the previous context was indexed by the surprisal of all the words in the naturalistic stimuli. Surprisal evoked by an incoming word indicates the amount of information that was not predictable from the context (Hale, 2001; Levy, 2008), and is calculated as the negative logarithm of the probability assigned to the actual next word. The probability of each word in the naturalistic stimuli given the previous words within the same sentence was derived from the pretrained GPT2-large model. This model uses a transformer architecture and has been shown to successfully capture human performance on next-word prediction (e.g., (Goldstein et al., 2022; Schrimpf et al., 2021)). The analyses was performed using the python package transformers (v4.10.2).

### Emotion arousal and emotional valence

Emotional arousal and emotional valence of each sentence in the naturalistic stimuli were rated by participants on Amazon Mechanical Turk (MTurk). Following a prior study (Wallentin et al., 2011), arousal was rated on a scale from 0 to 10 indicating extreme boredom to extreme arousal. Emotional valence was rated on a scale from −5 to 5 where −5 indicates strong negative emotions and 5 indicates strong positive emotions. A total of 30 participants completed the survey. The mean valence and arousal ratings for each sentence were computed, and words in the same sentence have the same emotional arousal and emotional valence. Inter-subject correlations (ISC) among each subject’s ratings for arousal and valence were computed as the mean of the Pearson’s *r* coefficients between each subject’s ratings and the overall mean ratings. The statistical significance of subjects’ ISC coefficients was determined by comparing the observed values with randomly generated ratings using paired two-sample *t*-tests.

## Results

### Behavioral results for the two-word task

Overall, participants achieved an accuracy of 94.4% (SD=23%) with a mean reaction time (RT) of 2.6 s (SD=2.07 s). The mean accuracy for adults was 96.9% (SD=17.3%) and the mean accuracy for children was 89.9% (SD=30.1%). The mean RT for adults was 2 s (SD=1.1 s) and the mean RT for children was 3.67 s (SD=2.83 s; see Figure 1B). Compared to prior studies (e.g., (Bemis & Pylkkanen, 2011)), these RTs seem longer. This is because the task was more difficult as the participants needed to use two buttons to select from 8 pictures. The reason for the more complex task was to reduce the possibility of correct responses by chance, which makes the task more applicable for possible clinical uses.

The binary variable Accuracy was analyzed using a generalized linear mixed-effects model (GLMM) with binomial error distribution, and RTs were log-transformed and analyzed using a linear mixed-effects model (LMM). Composition (single nouns vs. phrases) and Age (adults vs. children) were included as fixed effects and subjects as random intercepts. The results revealed a significant effect of Age on both accuracy (*p*<.001) and RT (*p*<.001). Composition was significant for RT (*p*=0.0003) but not accuracy (*p*=0.94).

### Phrasal and noun representation in LLMs

To gain insights into the neural representations of phrases and single nouns in the two-word and naturalistic contexts, we first examined the pretrained GPT2-large model’s embeddings of adjective-noun phrases and single nouns in a two-word setting (e.g., “green glass” vs. “glass”) and a naturalistic setting (e.g., “…soft music…” vs. “…a bath…”). The MDS results showed that in the two-word context, there is a clear separation of noun and phrasal representations in the LLM. In the naturalistic setting, however, the last layer’s representations of nouns and phrases were both highly distributed (see Figure 2A). The cosine distances between each layer’s embeddings of adjective-noun phrases and single nouns in the two contexts were shown in Figure 2B. We can see a larger distance in the middle and final layers of the LLM.

### Classification results on LLM embeddings

To understand whether the LLM has learned the contrast between single nouns and adjective-noun phrases, we trained two feed-forward neural network classifiers to distinguish phrases from nouns in the two-word context and tested the trained classifiers in the naturalistic context, and vice versa (see Figure 2C). The classifier trained in the two-word context achieved an accuracy of 80% in distinguishing phrases from nouns and an accuracy of 60.8% when applied to the naturalistic context. The classifier trained in the naturalistic context achieved an accuracy of 83.3% and an accuracy of 64.3% when tested in the experimental context (see Figure 2D). Although the testing accuracies were much lower than the training accuracy, the results in the two-word and naturalistic settings were comparable and were well above the chance level of 50%, suggesting that the LLM has learned different representations for single nouns and adjective-noun phrases and can be generalized across contexts.

### Classification results on MEG data

We applied the same classification methods to the MEG data to examine the generalizability of the neural reflections of semantic composition. Figure 3B shows the classification results of adults’ and children’s MEG data. For adults, the classifiers trained on the experimental data can distinguish phrases from single nouns in the left anterior and middle temporal lobe from 240-320 ms (*p*=0.025) after the onset of the target word. The classifiers trained on the naturalistic data can distinguish phrases from single nouns in the similar left anterior and middle temporal regions from 560-680 ms (*p*=0.03) after the onset of the target word. For children, the classifiers trained on the experimental data can distinguish phrases from single nouns in the left middle temporal lobe from 300-420 ms (*p*=0.014) after the onset of the target word. The classifiers trained on the naturalistic data can distinguish phrases from single nouns in the left anterior and middle temporal regions from 560-640 ms (*p*=0.001) after the onset of the target word. Since the adults’ children’s results exhibited similar spatiotemporal patterns, we collapsed their data together for future analyses.

For all subjects’ data, we found that when trained on the two-word data, the classifiers can distinguish phrases from single nouns in the left anterior and middle temporal lobe from 200-340 ms (*p*=0.0052) after the onset of the second word. When tested on the naturalistic data, the TGM results suggest that the classifiers from 200-220 ms in the training data can significantly distinguish phrases from nouns from 700-760 ms in the testing data. When trained on the naturalistic data, the classifiers can distinguish phrases from single nouns in the whole left temporal lobe from 520-680 ms (*p*=0.001) after the onset of the word. When tested on the experimental data using TGM, the classifiers from 620-640 ms in the training data can significantly distinguish phrases from nouns from 220-340 ms in the testing data (see Figure 3C).

### Neural dynamics of phrasal and noun representations

We used MDS to visualize the neural codes associated with each adjective-noun phrase and single noun in the two-word and naturalistic contexts. Within the significant spatiotemporal clusters derived from the classification analyses, we plotted the averaged MEG data of each phrase and noun in a 2-dimensional space. We also plotted the temporal dynamics of the mean 2D neural codes for all phrases and nouns in the two contexts. The results suggested reliable segregation of multivariate neural signals associated with adjective-noun phrases and single nouns in both experimental and naturalistic contexts. However, the temporal dynamics of the MDS representations also showed different patterns in the two contexts: In the two-word setting, the neural distance between phrases and nouns was larger in an earlier time window at around 100-400 ms and converged near the end of 800 ms. In the naturalistic setting, the neural codes for phrases and nouns remained distant from around 400 ms to the end of the epoch (see Figure 4). This is consistent with the classification results where the composition effect occurred later in the naturalistic context.

### Regression model of the naturalistic MEG data

Figure 5B shows the distributions of these regressors for the naturalistic stimuli. The mean root-mean-squared (RMS) intensity and mean f0 for all target words in the naturalistic stimuli were 0.21 A (SD=0.08 A) and 317.9 Hz (SD=40.02 Hz), respectively. The mean presentation rate of the target words in the naturalistic stimuli, calculated as the duration of the word divided by the number of letters in the word, was 72.8 ms (SD=23.5 ms). The mean log frequency and surprisal of the target words based on GPT2 in the naturalistic stimuli were 18.04 (SD=1.83) and 14.04 (SD=2.66). The mean emotional valence and arousal indicated whether the sentences containing the target words induced positive or negative emotion (−5 is very negative and 5 is very positive), and how strong the emotion was (on a scale of 0-10). Their mean values were 1.83 (SD=2.65) and 6.48 (SD=2.66). The mean intersubject correlations (ISC) among the participants’ ratings on valence and arousal were 0.74 (SD=0.13) and 0.57 (SD=0.2) and were both significantly greater than randomly generated ratings (*t*=20.24, *p*<.0001 and *t*=13.66, *p*<.0001, respectively), suggesting high agreement among the subjects on the two emotional dimensions associated with sentences in the naturalistic stimuli. We also examined the correlation coefficients among the regressors and the results suggested no collinearity among the regressors. The correlation coefficient between emotional valence and emotional arousal is −0.3, which is the highest absolute *r* value among all the regressor pairs (see Figure 5C).

### Classification results of the naturalistic data after regressing out other factors

We took the residuals of the source estimates for the target words in the naturalistic stimuli for each subject, and re-conducted the same classification analyses on the residuals. Our results confirmed that the late composition effect observed in the naturalistic data was due to additional processing efforts of these factors: The classifiers trained on the naturalistic data distinguished phrases from single nouns in a large cluster in the left temporal lobe from 280-400 ms (*p*=0.002) after the onset of the word. When tested on the experimental data using TGM, the classifiers from 320-340 ms in the training data significantly distinguished phrases from nouns from 220-360 ms in the testing data (see Figure 5D).

## Discussion

Traditional experimental paradigms in cognitive neuroscience of language aim to isolate specific cognitive processes by comparing conditions that differ in the component of interest. In contrast, recent naturalistic paradigms use audiobooks or movies to mimic everyday language experiences. However, both paradigms have limitations. Controlled experimental stimuli may deviate from natural language use, and subtraction methods assume linearity in a brain that is likely non-linear (Friston et al., 1996). Naturalistic stimuli contain diverse linguistic and non-linguistic information, making it challenging to isolate specific subprocesses (Hasson & Egidi, 2015). Direct comparisons of neural signals for linguistic processes between the two paradigms are rare, leaving it unclear if results from traditional experiments generalize to naturalistic settings and vice versa. According to existing neurolinguistic models (e.g., Hickok & Poeppel, 2000) brain areas associated with specific functions should not vary with the research context. For example, the left anterior temporal regions’ involvement in semantic composition should be consistent during phrasal processing, regardless of the paradigm or modality of stimuli presentation.

This study investigates the generalizability of meaning composition across traditional experimental and naturalistic paradigms, focusing on the core function of human language. We examined whether semantic composition observed in experimental paradigms extends to a naturalistic setting, and vice versa. Our classification results revealed similar neural activity for meaning composition in the left anterior and middle temporal regions in both experimental and naturalistic contexts. These findings align with previous studies employing similar two-word designs (e.g., Blanco-Elorrieta et al., 2018; Li & Pylkkänen, 2021; Westerlund & Pylkkänen, 2014, reviewed in Pylkkänen, 2019). Notably, the classification performance extended beyond the significant clusters observed in the two-word data, indicating the involvement of a larger network during the naturalistic task (as depicted in Figure 3B, first column). One line of research suggests that there is a hierarchy of increasing temporal receptive windows from lower sensory to higher perceptual and cognitive brain areas, and different levels of linguistic units are encoded at different cortical regions (e.g., Blank & Fedorenko, 2020; Hasson et al., 2008; Lerner et al., 2011; Schmitt et al., 2021). It is possible that phrasal processing in the naturalistic context encompasses longer temporal receptive windows, considering the richer contextual information, thus engaging more anterior or posterior temporal regions compared to isolated phrases.

Consistent with the hypothesis of longer temporal receptive windows, our findings revealed a delayed distinction between single nouns and adjective-noun phrases in the naturalistic MEG data, occurring from 520-680 ms after the onset of the target word, compared to the effect observed in the two-word task from 200-340 ms. Both our Temporal Generalization Mapping (TGM) and Multidimensional Scaling (MDS) analyses on the MEG data supported this latency contrast for composition in both paradigms. As naturalistic stimuli encompass richer information, including diverse prosodic cues, word rate, word frequency, and surprisal evoked by incoming words (Hale, 2001; Levy, 2008), as well as non-linguistic factors like emotional arousal and valence (Wallentin et al., 2011), prior neurolinguistic studies employing a naturalistic design have commonly controlled for these factors using regression models (e.g., Brennan et al., 2016; Caucheteux & King, 2022; Huth et al., 2016; Nelson et al., 2017). In our study, we accounted for these factors by regressing them out and then conducted the classification analyses using the residuals. Interestingly, after controlling for these factors, we observed an earlier composition effect that closely resembled the effect observed in the two-word data. This suggests that the composition effect observed in both experimental and naturalistic approaches reflects the same underlying processes, rather than being distinct processes.

Similarly, the classification results on the embeddings of single nouns and adjective-noun phrases in both the two-word and narrative contexts of the large language models (LLMs) indicate the presence of generalized patterns for these word types. While the question of whether these patterns reflect composed meaning in LLMs remains open, the results demonstrate the existence of specific features that differentiate single nouns from adjective-noun phrases and can be generalized across different contexts. It is important to note that the two-word stimuli introduce a confounding factor, as the nouns in the single nouns condition are the initial tokens, while the nouns in the adjective-noun phrases condition are the second tokens. To mitigate this factor, we deliberately removed the word position effect from each layer’s embeddings, ensuring that the classifier cannot rely solely on word position to distinguish between the two conditions.

To sum up, we observed the composition effect in both the experimental and naturalistic designs in similar brain regions and similar temporal windows when controlled for additional factors in the naturalistic stimuli, suggesting a single compositional process during both isolated and connected speech comprehension. One limitation of our study is that we only focused on a specific linguistic subprocess, and further research is needed to examine whether other subprocesses, such as morphological or syntactic processing, can be replicated across different research paradigms. Conducting meta-analyses using existing experimental and naturalistic fMRI datasets from open data platforms could serve as a valuable starting point for future investigations in this direction.

## Data and code availability

The data and the codes for analyses are available at https://osf.io/7c58j/.

